# Conserved gene content and unique phylogenetic history characterize the ‘bloopergene’ underlying *Triturus’* balanced lethal system

**DOI:** 10.1101/2024.10.25.620277

**Authors:** Manon Chantal de Visser, James France, Olga Paulouskaya, Thomas Brown, Michael Fahrbach, Chris van der Ploeg, Ben Wielstra

## Abstract

In a balanced lethal system, half the reproductive output succumbs. *Triturus* newts are the best-known example. Their chromosome 1 comes in two distinct versions and embryos carrying the same version twice experience developmental arrest. Those possessing two different versions survive, suggesting that each version carries something uniquely vital. With target capture we obtain over 7,000 nuclear DNA markers across the genus *Triturus* and all main lineages of Salamandridae (the family to which *Triturus* belongs) to investigate the evolutionary history of *Triturus’* chromosome 1 versus other chromosomes. Dozens of genes are completely missing from either one or the other version of chromosome 1 in *Triturus*. Furthermore, the unique gene content of 1A versus 1B is remarkably similar across *Triturus* species, suggesting that the balanced lethal system evolved before *Triturus* radiated. The tree topology of chromosome 1 differs from the rest of the genome, presumably due to pervasive, ancient hybridization between *Triturus’* ancestor and other newt lineages. Our findings accentuate the complex nature of *Triturus’* chromosome 1 – the ‘bloopergene’ driving the evolutionarily enigmatic balanced lethal system.

## Introduction

A balanced lethal system is an extreme hereditary disease that, in diploid organisms, causes the loss of fifty percent of offspring every generation^1^. A balanced lethal system is characterized by the presence of two versions of a specific chromosome that do not recombine and that each harbor one or more unique, recessive lethal alleles^1, 2^. As a consequence, individuals heterokaryotypic for this chromosome (i.e., individuals that inherited both versions), possess all genes required for survival^1, 3^. However, given the laws of Mendelian inheritance, half the offspring in each generation comprise individuals that are homokaryotypic for the chromosome, meaning they possess either one or the other version twice^4, 5^. Such individuals miss certain crucial genes, causing them to be inviable. It seems an evolutionary paradox that, despite the huge loss of fitness incurred, balanced lethal systems have evolved repeatedly across the tree of life^2, 5, 6, 7, 8, 9^.

Given that the two chromosome versions involved in a balanced lethal system do not recombine, they should be considered supergenes: physically linked set of genes that are inherited together^10, 11, 12^. Different versions of a supergene generally encode complex, distinct phenotypes that are maintained by balancing selection when each offers a unique advantage^13, 14^. However, suppressed recombination in supergenes – often caused by structural variation such as chromosomal inversions^10^ – also comes with drawbacks. Because purifying selection is weakened under reduced recombination, supergenes tend to accumulate deleterious alleles and recessive lethality may develop in homokaryotypes over time^1, 3^. For instance, this has been proposed to have caused the evolution of the recessive lethal mutations embedded in the supergenes found in fire ants^15, 16^. Moreover, supergene-bound recessive lethal genes may originate suddenly, due to a disruptive inversion breakpoint. For example, such a breakpoint mutation affects the inversion-based supergene that underlies the complex male mating strategy polymorphism in ruffs and is lethal in the homozygous state^17, 18^. Thus, a supergene might instantly qualify as, or gradually turn into, what we call a ‘bloopergene’: a genetic construction that can boost the rapid evolution of adaptive traits, but likewise is an evolutionary trap because of maladaptive consequences. Following this rationale, a balanced lethal system should be considered the most extreme case of a ‘bloopergene’.

The best-known example of a balanced lethal system concerns ‘chromosome 1 syndrome’ in the salamander genus *Triturus*: the crested and marbled newts^1, 19, 20^. Over two centuries ago, Rusconi already observed that half of *Triturus* embryos perish while still inside the egg^21^. From karyotyping studies in the 1980s it became clear that the fundamental cause of this massive die-off is a balanced lethal system^20, 22^. *Triturus* newts possess two versions of chromosome 1, dubbed 1A and 1B, that lack chiasmata and do not recombine along most of the long arm^20^. Only heterokaryotypic (1A1B) individuals survive the balanced lethal system, while homokaryotypic individuals (1A1A and 1B1B) experience developmental arrest^19, 23, 24, 25^.

All *Triturus* species share chromosome 1 syndrome, indicating that this balanced lethal system originated in their common ancestor and has, remarkably, persisted for at least 24 million years^1, 26^. On the one hand, a block-like deletion structure is known to characterize the 1A- and 1B-specific regions in the *Triturus* linkage map, suggesting a rapid origin due to a chromosomal re-arrangement – a ‘cytogenetic accident’^24, 27^. On the other hand, different *Triturus* species exhibit different Giemsa C-banding patterns in both 1A and 1B, suggesting that 1A and 1B have accumulated differences in genomic content between species after the balanced lethal system became fixed^25, 28^. Thus, determining the degree of any genetic divergence across *Triturus* is required to reconstruct the ancestral constitution of chromosome 1 and help us better understand the evolutionary origin of the balanced lethal system.

Supergenes are known to arise occasionally through introgressive hybridization^29, 30^ and such a scenario could theoretically kickstart a balanced lethal system by bringing together two distinct versions of a chromosome, each ‘pre-loaded’ with private deleterious mutations, in a single population^3^. Salamandridae, the salamander family to which *Triturus* belongs, is known to have a complex evolutionary history and bears an extensive signal of introgressive hybridization^31, 32, 33^. While the ‘cytogenetic accident’ hypothesis would predict that 1A and 1B originated in a single ancestral *Triturus* population^27^, the possibility that 1A and 1B were brought together into a single population^3^ by introgressive hybridization should not be dismissed out of hand. To test these hypotheses we can compare the evolutionary history of *Triturus*’ chromosome 1A and 1B to that of the other chromosomes.

Here, we conduct a genome-wide investigation using target capture data of 148 samples from 16 genera within the Salamandridae family (covering 26 species in total), with the aim of unraveling the evolution of *Triturus*’ peculiar chromosome 1. Firstly, we identify presence/absence variation associated with the balanced lethal system in *Triturus* by comparing 1A- and 1B-linked markers across ten different *Triturus* species to explore how the gene content varies. Secondly, we investigate the evolutionary history of *Triturus’* chromosome 1 versus that of the rest of the *Triturus* genome by conducting phylogenomic analyses including all main Salamandridae lineages. We interpret our results in the light of existing and new theories regarding the origin of balanced lethal systems.

## Results

### Consistent deletions on 1A and 1B across *Triturus*

To test the hypothesis that the 1A- and 1B-specific regions of different *Triturus* species vary in genomic content, we use target enrichment by sequence capture to aim to obtain 7,139 nuclear DNA markers^34, 35^ from 1A1A, 1A1B and 1B1B embryos belonging to ten *Triturus* species (n=3 for each genotype, for each species, as listed in Table S1). In healthy (1A1B) embryos, 6,785 genes are recovered. We explore which of these genes are missing in either 1A1A embryos (and therefore presumably absent from chromosome 1A) or 1B1B embryos (and therefore potentially absent from chromosome 1B) for each *Triturus* species. To strengthen our confidence that genes are truly absent from 1A or 1B, we determine their position on linkage maps of *Triturus* and its sister genus *Lissotriton;* the smooth newts^27^, the chromosome-scale assembly of the distantly related *Pleurodeles;* the sharp-ribbed newt^36^, and an Oxford Nanopore sequencing-based *Triturus* assembly, scaffolded against the *Pleurodeles* genome, with the understanding that synteny in newts is high^27^. We find that 29 genes are consistently absent *Triturus*-wide in diseased 1A1A embryos (but present in 1A1B and 1B1B embryos; Fig.1 and Table S2) and another 35 genes are consistently absent across *Triturus* in diseased 1B1B embryos (but present in 1A1B and 1A1A embryos; Fig.1 and Table S2).

**Fig 1.**
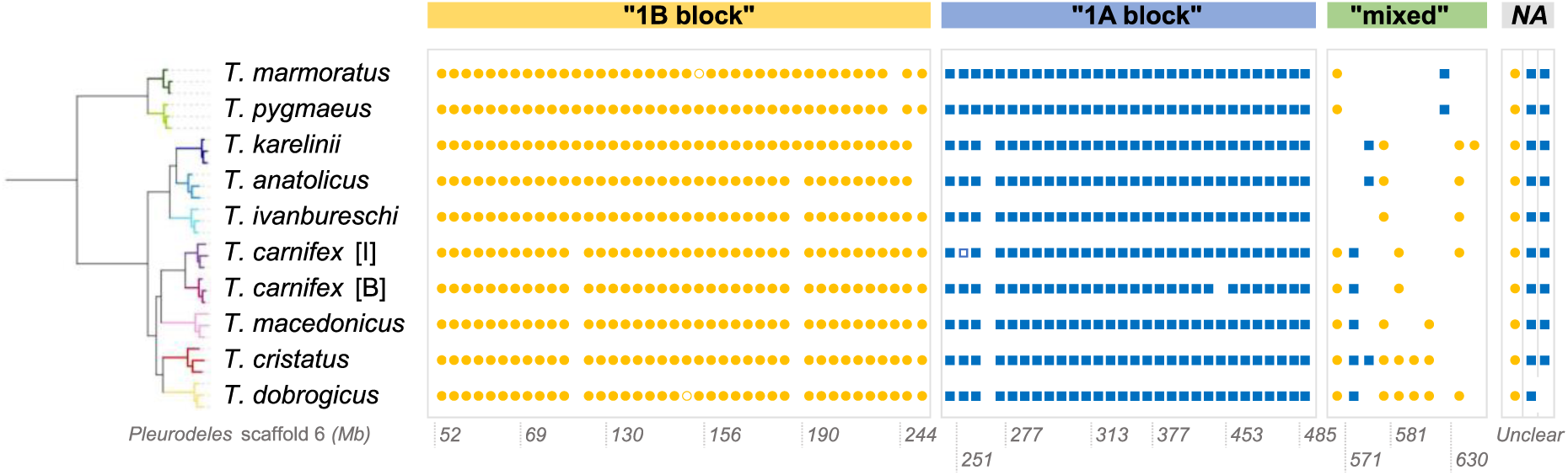
Presence/absence variation matrix showing variation in 1A- and 1B-linked markers across the genus Triturus. For each Triturus species, three individuals with the 1A1A genotype, three with the 1A1B genotype, and three with the 1B1B genotype, were compared. Each column in the matrix represents a recovered 1A-linked (blue square) or 1B-linked marker (orange circle). A gap indicates no presence/absence was detected and an open symbol reflects uncertainty. The matrix is ordered based on the position of the markers in the Pleurodeles genome (scaffold 6; location in Mbp indicated at the bottom). From left to right there is a distinct 1B-block containing 40 markers, followed by a distinct 1A-block containing 30 markers, followed by a relatively small ‘mixed’ block with three 1A- and seven 1B-linked markers. Three additional markers could not be mapped in Pleurodeles (‘Unclear’). A total of 35 1B-linked and 28 1A-linked markers are consistently found across all Triturus species. The Triturus phylogeny is based on 19,827 SNPs from 3,728 targets that are not linked to chromosome 1 (see also Fig. S35). [I] = Italy clade, [B] = Balkan clade.

The 1A and 1B chromosomes are remarkably similar in the different *Triturus* species in terms of gene content: variation in gene-deletions is low between species, and there is no apparent phylogenetic signal for the few gene-deletions that do deviate between species (Fig. 1). This indicates that deletions mainly happened before the radiation of *Triturus* and that only a few genes were lost afterwards. Also, the gene order along the chromosome-scale genome assembly of *Pleurodeles waltl*^36^, suggests that chromosomes 1A and 1B are characterized by unique, big deletions of DNA stretches that contain dozens of genes and together span ∼600 Mbp (in *Pleurodeles*). The genes deleted from 1A- and 1B generally form two consecutive clusters, followed by a small ‘mixed’ block consisting of both presence/absence variation markers, as well as markers with distinct copies on A and B (Fig.1 and Table S2). This ‘mixed’ block suggests the presence of a relatively small, evolutionary stratum present on both 1A and 1B, adjacent to the 1A-block in the estimated, ancestral constitution of chromosome 1.

Given the conserved synteny across different, distantly related newt genera^27^, and given our observation of little variation in 1A- and 1B-linked markers across the entire genus *Triturus*, we can now further explore the evolutionary history of chromosomes 1A and 1B compared to the rest of the *Triturus* genome.

### Discordant evolutionary history for *Triturus’* chromosome 1

To test the hypothesis that chromosomes 1A and 1B originated in different evolutionary lineages of newt and were later brought together in a single population through introgressive hybridization, we sequence our c. 7k nuclear DNA markers for all main evolutionary lineages in the Salamandridae family (Table S1). We then build phylogenetic trees for markers from all twelve linkage groups (presumed to correspond to the twelve chromosomes) of *Triturus* separately^27^, and we also build trees for only the recombining section of chromosome 1 (i.e. predominantly the short arm), the 1A-linked markers, and the 1B-linked markers, to explore whether either the 1A- or the 1B-linked markers display an aberrant evolutionary history.

We observe phylogenetic discordance in the ‘modern European newts’, a group of newts composed of four main lineages: the genera *Triturus*, *Lissotriton* and *Calotriton*, as well as the ‘NIO clade’, which comprises the genera *Neurergus*, *Ichthyosaura* and *Ommatotriton*^33^. The topology for the entire chromosome 1, rather than only the 1A-specific or 1B-specific regions, deviates from that of the rest of the genome (Fig. 2 and Figs. S1-S18). Individual phylogenies for linkage groups 2 through 12 suggest that *Lissotriton* is the sister lineage of *Triturus* and that, together, these two genera comprise the sister lineage of the NIO clade. On the other hand, both the recombining section of chromosome 1, as well as the separate 1A- and 1B-linked markers, suggest that the phylogenetic positions of the NIO clade and *Lissotriton* are switched, with the NIO clade recovered as the sister lineage of *Triturus* (although for 1B-linked markers the support for *Triturus* + the NIO clade is relatively low). The 1B-linked markers show further deviation still: not *Lissotriton,* but *Calotriton* (more distantly related for all the other linkage groups) is suggested to be sister to the *Triturus* + the NIO clade, albeit with very low support (Fig. 2). Lastly, molecular dating suggests that the radiation of all newt clades involved (thus *Triturus*, *Lissotriton*, the NIO clade, and *Calotriton*, together known as the ‘modern European newts’) occurred in the distant past and in a brief time window (c. 38-42 million years ago; Figs. S18-S34). Given this topological discordance, we investigate the genomic pattern of introgression in more detail.

**Fig 2.**
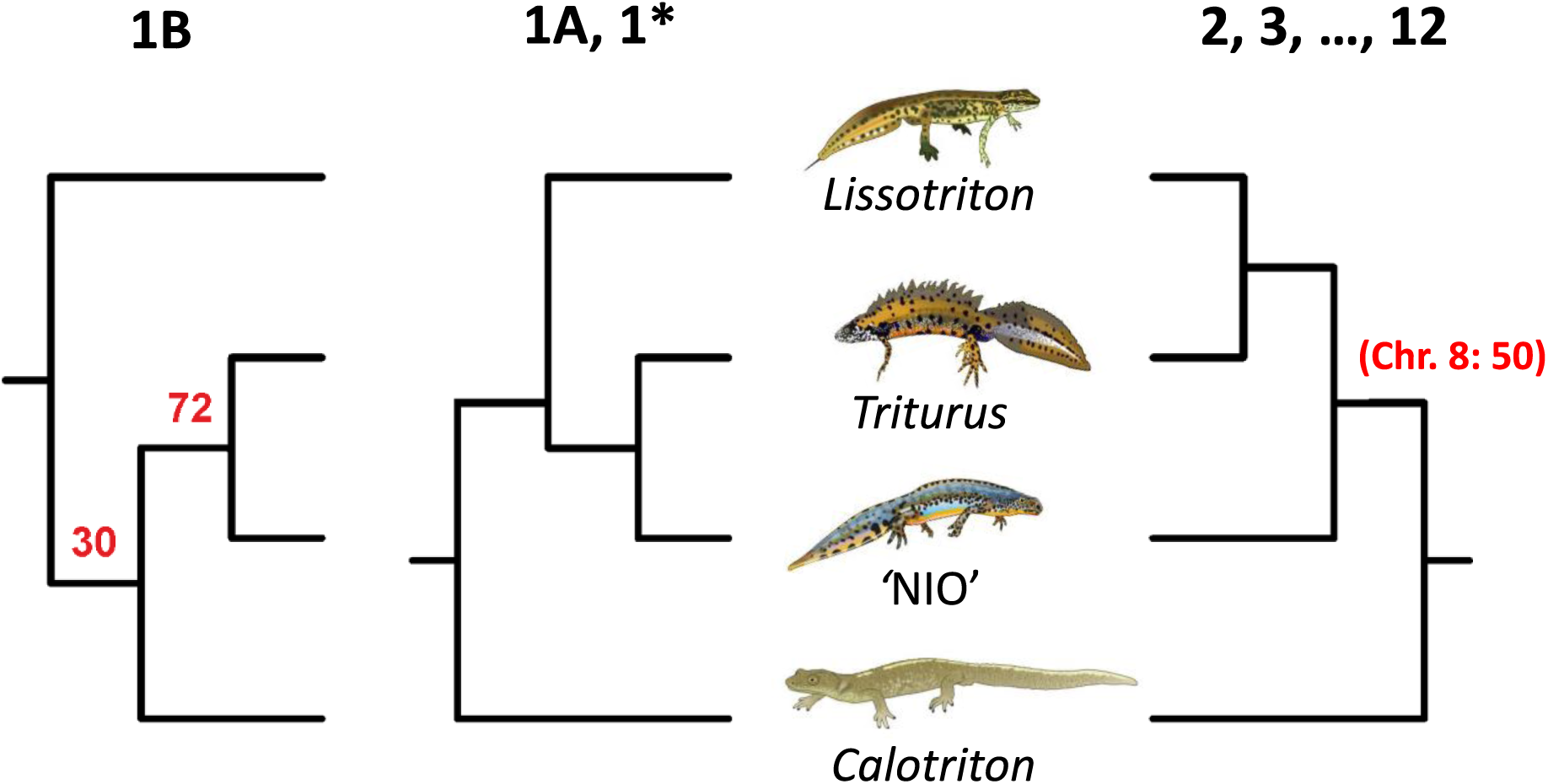
*Topologies for the four clades that comprise the ‘modern European newts’ for different parts of the genome. The topology on the far right (2-12) shows the result for targets of chromosomes 2 through 12 combined, as well as for each of the 12 linkage groups separately. The two topologies on the right show the result for the recombining region of chromosome 1 (1*), for chromosomes 1A (same as 1*), and for chromosome 1B – together comprising linkage group 1. Red numbers indicate three cases where the bootstrap support was below 80 (see Figs. S1-S18). The NIO clade is a consistently recovered monophyletic group that contains the three genera* Neurergus, Ichthyosaura *and* Ommatotriton.

### A complex history of pervasive introgression in modern European newts

The discordant phylogenetic position of *Triturus* within the modern European newts, based on chromosome 1 versus the rest of the genome, implicates a history of introgression. We test for deviations from a strictly bifurcating evolutionary history in order to identify potential key introgression events, again for markers positioned on the different *Triturus* linkage groups, including 1A, 1B and the recombining section of chromosome 1. Our analyses show that a signal of introgression involving the NIO clade affects not only *Triturus’* chromosome 1, but most other linkage groups as well (Fig. 3). To a lesser extent, introgression involving *Calotriton* is also implicated, which is in line with the observation that mtDNA suggests this taxon to be the sister lineage of *Triturus*^33^. Overall, the dominant signal of introgressive hybridization suggests introgression from the ancestor of the NIO clade into the ancestor of the *Triturus* lineage (Fig. 4).

**Fig 3.**
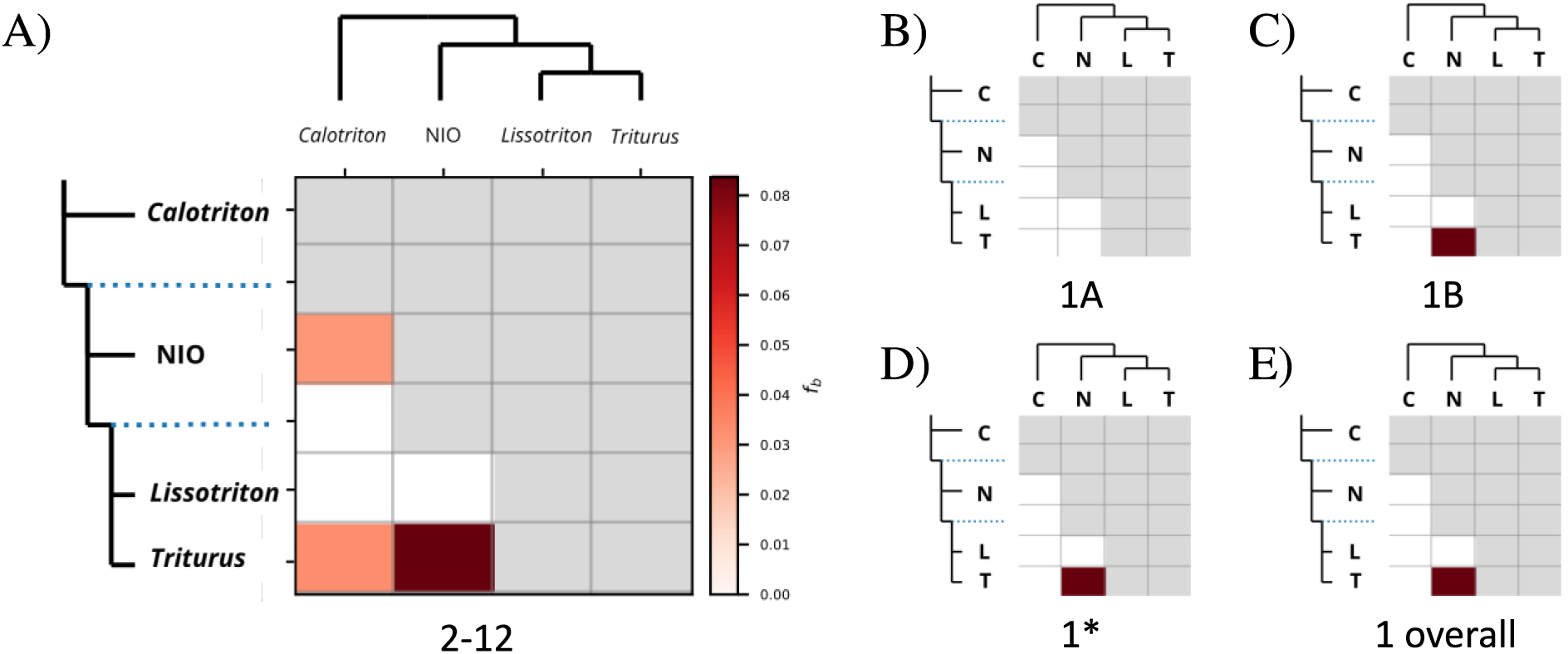
*F-branch (f_b_) heatmaps for different subsets of markers/linkage group(s), showing signals of pervasive introgression in the past among ‘modern European newts’. Colour shading reflects the intensity of excess allele-sharing (indicative of introgression) between tree branches on the y-and x-axes. Grey boxes reflect it is not possible to calculate f-branch statistics following the tree topology. Blue, dotted lines correspond to the internal tree branches (e.g. the line above* Lissotriton *and* Triturus *represents their common ancestor). (A) Heatmap based on markers positioned on* Triturus *linkage groups 2 through 12 (n=3,728 targets). (B, C, D, E) Summarized heatmaps for, respectively; 1A-linked markers only (n=28 targets), 1B-linked markers only (n=35 targets), markers positioned on the recombining region of chromosome 1 (1*, n=329 targets), and all markers found on chromosome 1 (n=390 targets). A noticeable, strong signal for introgression between* Triturus *and the NIO clade is visible, except in the analysis including 1A-linked markers only (which notably has the lowest sample size). Abbreviations in figure B-E: T =* Triturus*, L =* Lissotriton*, N = the NIO clade (a consistently recovered monophyletic group that contains the three genera* Neurergus, Ichthyosaura *and* Ommatotriton*), and C =* Calotriton*. See Table S3 and Figs. S36 – S86 for all heatmaps produced*.

**Fig 4.**
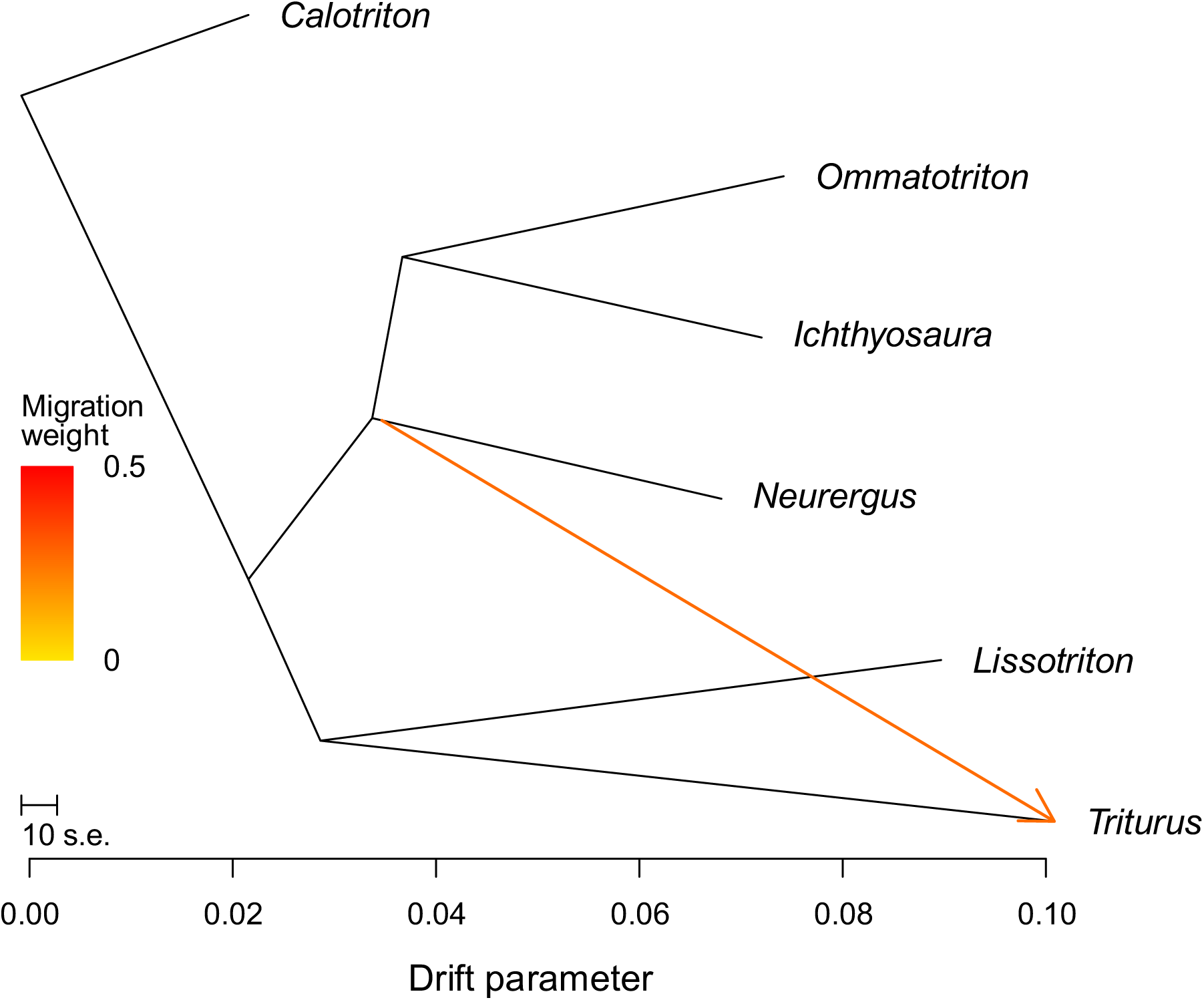
TreeMix admixture graph for the ‘modern European newts’ with one migration edge, based on the overall target capture dataset. The arrow suggests that introgression occurred between the ancestors of the NIO clade and the Triturus genus and shows the inferred direction of the migration event. The colour of the arrow indicates the migration weight. The axis at the bottom shows the genetic drift measured in the populations.

## Discussion

The chromosome underlying the balanced lethal system in *Triturus* has a remarkable evolutionary history. Gene-content-wise, little has changed in 1A and 1B since the balanced lethal system became established in the *Triturus* ancestor. This is surprising, because natural selection would be expected to be less efficient at preventing the further shedding of genes since the system became fixed, as homokaryotypes from that point on would have been destined to die anyway^1, 3, 37^.

We see that the 1A- and 1B-specific regions together span ∼600 Mbp in the genome of *Pleurodeles*^36^, which is considerably smaller than would be expected based on the physical size inferred in karyotyping studies of *Triturus*, estimated to be at least 1.2 Gb^25^ – over a third of the size of the human genome. The affected region includes a small, evolutionary stratum that suggests restriction of recombination extends beyond the core ‘bloopergene’, allowing further divergence to have accumulated between 1A and 1B (although the pattern is notably minor). The variation in Giemsa C-banding patterns that is observed between different lineages of the genus presumably reflects the independent accumulation of non-coding DNA^25, 28^. This is not unexpected, given that salamander genomes are dominated by transposable elements, which tend to accumulate in non-recombining regions with weak, or absent, selection pressures^38, 39, 40^. However, even though the non-recombining long arms of chromosome 1 – comprising 1A and 1B – are physically large, the gene density is evidently low.

The near-identical gene content of 1A and 1B across the genus *Triturus* (in combination with the lack of a phylogenetic signal for the minor variation between species that is observed), supports that the ‘bloopergene’ driving the *Triturus* balanced lethal system originated *before* the onset of the radiation of *Triturus* occurred. This fits the hypothesis that chromosomes 1A and 1B evolved instantaneously due to a ‘cytogenetic accident’^24, 27^. But does that preclude that introgressive hybridization played a role in the origin of the balanced lethal system?

Our phylogenomic approach reveals a divergent topology for chromosome 1 compared to the rest of the *Triturus* genome. The entire *Triturus* genome bears the signature of introgression derived from the NIO clade. However, introgressive hybridization seems to have affected chromosome 1 in particular, to the extent that a sister relationship between *Triturus* and the NIO clade is supported in our phylogenetic analysis (rather than *Lissotriton* being the sister lineage of *Triturus* as is the case for the rest of the genome). Yet, the scenario that chromosome 1 reflects the actual evolutionary relationships and was relatively resistant to introgression from *Lissotriton* compared to the rest of the genome can perhaps not be dismissed out of hand, given that including more molecular markers does not strictly lead to a more accurate species tree^41, 42, 43^.

Our findings do not fit the hypothesis that introgression brought together supergenes that evolved in two different ancestral populations^3^. The confounding topology was observed not only for 1A-specific regions or for 1B-specific regions of chromosome 1, but for both of them – as well as for the rest of chromosome 1. This further supports that the balanced lethal system arose strictly within the ancestral *Triturus* population – in line with the hypothesis that chromosomes 1A and 1B originated in a cytogenetic accident^24, 27^. Still, the aberrant evolution of chromosome 1 in *Triturus* evidently did involve introgressive hybridization.

In conclusion, we reconstruct the ancestral constitution of *Triturus’* chromosome 1 and provide new insights into the evolutionary mechanisms that twisted the *Triturus* genome in the distant past. The ‘bloopergene’ that drives the balanced lethal system evolved over 24 million years ago within the ancestor of *Triturus*, against a backdrop of pervasive, introgressive hybridization, involving the other lineages that gave rise to the modern European newts. By unveiling the intricate genomic architecture and multifaceted history of the *Triturus* balanced lethal system, our results – paradoxically, much like the phenomenon itself – both enhance and challenge our understanding of how supergenes and ‘bloopergenes’ exactly originate and evolve in nature.

## Methods

### Sampling scheme and phenotypic + genotypic classification

To investigate the intergeneric variation of *Triturus*’ chromosome 1A and 1B, we obtain embryos from captive populations of all *Triturus* species listed in Table S1^44^. We include both the Balkan and Italian clade of *T. carnifex*, which are genetically distinct and presumably represent distinct species^45^. We do not include *T. rudolfi*^46^, closely related to *T. pygmaeus*, as it was only described after our sampling. Individuals were (pure)bred for an unknown number of generations in captivity, but the original founders of all breeding lines are known to derive from localities away from any (potential) hybrid zones^47^, ensuring interspecific admixture is limited. Husbandry and breeding practices are described in^48^.

The two types of homokaryotypic *Triturus* embryos (1A1A and 1B1B) express different phenotypes upon developmental arrest: the so-called “slim-tailed” (1A1A) embryos appear morphologically similar to healthy heterokaryotypic (1A1B) embryos, whereas the so-called “fat-tailed” embryos (1B1B) show more severe malformations^24, 25, 49^. During the sampling of *Triturus* embryos, embryos are thus classified as being healthy when they developed beyond the critical late tailbud stage, or as being diseased (in one, or the other, phenotypic group) when they experienced developmental arrest during this critical stage. In addition, we confirm this embryonic classification by genotyping samples using a 1A-linked marker (PLEKHM1), a 1B-linked marker (NAGLU), and a control marker from elsewhere in the genome (CDK) with multiplex PCR^50, 51^. In total, we include 90 *Triturus* embryo samples: three individuals per 1A1B, 1A1A, and 1B1B genotype, for ten species/populations.

For building a *Triturus* species tree, we use previously published data of 27 adult *Triturus* newts^34, 52^ and add three *T. carnifex* samples^45^ to account for presumed species-level divergence within this taxon (Table S1). Furthermore, to determine the phylogenetic relationships of 1A and 1B compared to the rest of the *Triturus* genome, we include other newts from the Salamandridae family in our study: we use three samples for closely related lineages (the modern European newts, comprising of the genera *Lissotriton, Ommatotriton, Ichthyosaura, Neurergus* and *Calotriton*) and at least one sample as representative for more distantly related lineages^33, 35^. This means that, next to the *Triturus* samples, we use 28 Salamandridae samples from 15 different genera (Table S1).

### Laboratory procedures and pre-processing of sequence data

To obtain genomic data we use “NewtCap”: a target enrichment by sequence capture workflow^35^. In brief, we apply a standard salt-based method for extracting DNA using the Promega Wizard^TM^ Genomic DNA Purification kit (Promega, Madison, WI, USA), followed by library preparation using the NEBNext Ultra^TM^ II FS DNA Library Preparation Kit for Illumina (New England Biolabs, Ipswich, MA, USA). Then, we perform target enrichment using a MyBaits-II kit using sequence-specific, *Triturus*-based RNA probes Arbor Bioscience, Ann Arbor, MI, USA, product Ref: # 170210-32^34, 35^ and Illumina sequencing of the enriched targets (outsourced to Baseclear B.V., Leiden, the Netherlands). An upstream bioinformatics pipeline is used to check the quality of the reads and to perform clean-up and mapping upstream^35^. Downstream analyses are described below.

### Oxford Nanopore sequencing

To gather as much information about the (likely) genomic position of markers within the DNA of *Triturus*, we perform whole genome sequencing, followed by *de novo* assembly. We use liver tissue from a *T. ivanbureshi* individual, flash frozen in liquid nitrogen and stored at −80°C prior to whole genome sequencing. High molecular weight DNA extraction and DNA sequencing using Nanopore is performed by Future Genomics Technologies. DNA is prepared using the SQK-LSK110 ligation library kit. FLO-MIN106 and FLO-PRO002 (R9.4.1) flow cells are used for sequencing on MinION and PromethION platforms. For basecalling, Guppy v.5.0.17^53^ is used with the high-accuracy model. From 14 flowcells, we obtain 91,494,016 reads, with an N50 ranging from 14,7 kb to 27,2 kb per flow cell. Using Porechop^54^ we trim adaptor sequences off of the raw reads.

### Genome assembly, scaffolding and alignment

We assemble the genome of *Triturus de novo* using the Oxford Nanopore Technology reads and scaffold the rather fragmentary output using the *Pleurodeles waltl* genome (56) as a reference to get a chromosome-level assembly. Although *P. waltl* does not suffer from a balanced lethal system, it is the most closely related genus with a published reference genome available and synteny with *Triturus* is high^27^, so it allows us to estimate the order of genes on *Triturus’* chromosome 1.

We run Shasta v.0.10.0^55^ with arguments: “--Reads.minReadLength 5000” and “--config Nanopore-May2022”, using reads > 5 kb, followed by removal of any retained haplotigs using purge-dups v.1.2.6^56^ to produce a draft *Triturus* genome assembly of 21 Gb in size. The assembly contains ∼65k contigs, with an N50 of 1.27 Mb. For scaffolding, we run RagTag^57^ with the *P. waltl* genome as input reference. This eventually scaffolds ∼17.6Gb of the 21Gb contigs assembly into the 12 separate chromosomes. We then use BLAST+ v.2.2.31^58^ to align the genes from the target capture set against our scaffolded genome assembly and *P. waltl* reference genome with default parameters. Only the genes that have one hit in each of the genomes are considered for further analysis.

### Presence/absence variation analysis with PAV-spotter

We use “PAV-spotter”^51^ to search for presence/absence variation in the target capture data of three diseased fat-tailed (1B1B), three diseased slim-tailed (1A1A), and three healthy/control (1A1B) embryos, for each *Triturus* species/population independently. PAV-spotter uses BAM file information to calculate signal-cross-correlation values that are based on the resemblance of the distribution of reads mapped against reference sequences for a given target. We use the following rationale: an absence of reads for a target across 1B1B samples is indicative of a 1A-linked marker in case reads are mapped for the same target across 1A1A and 1A1B samples, and likewise an absence of reads for a certain target across 1A1A samples denotes a 1B-linked marker when reads are mapped for the same target across 1B1B and 1A1B samples.

We prepare the read depth data for PAV-spotter and we apply the tool using default settings^51^. We use three samples per phenotypic category to correct for any stochasticity and variation in capture-rates between samples. To identify chromosome 1-linked presence/absence variation, we used a 80% dissimilarity threshold^51^. We confirm the presence/absence variation patterns by visually inspecting the read content in the BAM files for targets that are identified as either 1A- or 1B-linked markers using IGV Integrative Genomics Viewer^59^ and we again do this in the same way as described in^51^.

### Reconstructing the ancestral constitution of chromosome 1

After obtaining the list of candidates for 1A-linked and 1B-linked markers, we cross-compare the results for the different *Triturus* species. To determine if targets are true positives, we cross-check the data with 1) a *Triturus* linkage map^27^, 2) a *Lissotriton* linkage map^27^, 3) the *Pleurodeles* genome assembly^36^, and 4) the *Triturus* RagTag assembly (which is based on the *Pleurodeles* genome). In case targets do not show up in expected regions within these comparative genomic datasets, they are excluded as false positives (approximately 70% of potential 1A- and 1B-linked markers concerns singletons, meaning they are only discovered in one species).

As synteny across newt genera is high, but not absolute^27^, we follow a rationale in the order of cross-comparisons that is based on the relatedness to *Triturus*, where data of more closely related genera overrule the data of more distantly related genera. If a potential 1A- or 1B-linked target is found in the region of the *Triturus* linkage map that corresponds to the region of interest on chromosome 1 (between ∼0–50 cM on linkage group 1), we consider it a true positive. Next, for any remaining target that is not present in the *Triturus* linkage map, we consider it a true positive if it is present on the relevant region of the *Lissotriton* linkage map (between ∼50–100 cM on linkage group 4). For any remaining target at this point, we perform a cross-check with the best blast hits to the less closely related genus *Pleurodeles*. In case the target is recovered with a best blast hit on *Pleurodeles’* scaffold 6 between ∼0–500 Mb (which corresponds to the region of interest in *Triturus*), we include it. Finally, for any remaining target we cross-check against our *Triturus* RagTag assembly. In case we observe that a target is on scaffold 6 of the RagTag assembly between ∼0–500 Mb (again, the region of interest in *Triturus*), we include it. Any remaining targets are excluded.

### Building the *Triturus* phylogeny

To help visualize potential phylogenetic signal in the presence/absence variation matrix, we build a *Triturus* phylogeny. We exclude the entirety of chromosome 1 by restricting the input for the analysis to the 3,728 targets that were placed on linkage groups 2-12 in the *Triturus* linkage map^27^. To extract only high quality SNPs, we apply quality filtering by removing heterozygote excess, as well as by extracting SNPs and discarding INDELs, applying stringent quality filtering options, and removing sites with more than 50% missing data as described in^35^. Next, we remove invariant sites using an ascertainment bias correction Python script (https://github.com/btmartin721/raxml_ascbias) and convert the filtered VCF to PHYLIP format using the vcf2phylip script, v.2.8^60^ to serve as input for RAxML. Then, we run RAxML v.8.2.12 with 100 rapid bootstrap replicates under the ASC_GTRGAMMA model and Lewis ascertainment correction to perform a concatenation analysis and obtain the best-scoring maximum likelihood tree^61^, following the methods of^35^. This phylogeny (Fig. 1 and Fig. S35) was based on in total 19,827 SNPs. We visualized the tree using FigTree v.1.4.4 and enhanced the coloured version in Figure 1 using iTOL v.6^62^.

### Estimating Hardy-Weinberg Equilibrium deviations

To search for genes that are potentially still present on both chromosome 1A and 1B across (most of the) *Triturus* species, we search for targets that deviate from Hardy-Weinberg Equilibrium and display excessive heterozygosity explicitly in 1A1B embryos (and not in 1A1A or 1B1B embryos). We create three multi-sample VCF files for *Triturus*; one with the 30 1A1B embryos, one with the 30 1A1A embryos, and one with the 30 1B1B embryos. Then, we use BCFtools v.1.18^63, 64^ to estimate Hardy-Weinberg and related statistics for every site in each of the VCF file. To take into account potential gene losses, or stochasticity in the data, we allow for a maximum of three individuals to have a missing genotype at any particular site per each VCF file, and we filter as such using the *--max-missing* function of VCFtools^65^.

From the data we extract the sites for which heterozygote excess was discovered in the 1A1B set (with a p-value below 0.05), but neither in the 1A1A, nor in the 1B1B set (i.e., the p-value was above 0.05 in both). As before (see “Presence/absence variation analysis with PAV- spotter”), we check whether the discovered targets fall in the (vicinity of the) heteromorphic *Triturus* chromosome 1 region. Finally, we again perform a visual inspection in IGV^59^ to determine if heterozygous SNPs are restricted to 1A1B embryos. Based on a p-value of 0.05, we discover only six targets that show significant heterozygote excess in explicitly 1A1B embryos, pass our IGV inspection, and appear in or near the region(s) of interest (Table S1).

### Phylogenetic analyses of Salamandridae

To investigate if the topology of chromosome 1 differs from that of the rest of the genome, we build a species tree of Salamandridae, following the same phylogenetic approaches as described above. We perform separate RAxML analyses based on SNPs from chromosome-level subsets of targets mapped to different *Triturus* linkage groups: “Chr1” (all 390 targets on *Triturus* linkage group 1, including 1A- and 1B-linked markers), “Chr1*” (a subset of 329 targets on *Triturus’* recombining part of linkage group 1, thus excluding 1A- or 1B-linked markers), “1A” (the subset of 28 markers that was 1A-linked for all *Triturus* species analyzed), “1B” (the subset of 35 markers that was 1B-linked for all *Triturus* species analyzed), “Chr2- 12” (all 3,728 targets on *Triturus* linkage groups 2-12), and “Chr2” – “Chr12” (subsets of targets for each individual linkage group; see Table S3). We time-calibrate the best-scoring ML trees discovered by RAxML using TreePL^66^ using to strict calibration points: 67.49 MYA, for the split between the “New World Newts” and “Old World Newts” and 24 MYA for the most recent common ancestor of *Triturus*^26^. The trees are visualized in FigTree v.1.4.4 and shown in Figs. S1-S34.

### Calculating D- and f-statistics

To test the hypothesis that chromosomes 1A and 1B originated in different evolutionary newt lineages and later ended up in a single population through introgressive hybridization, we calculate Patterson’s D, also known as the ‘ABBA-BABA statistic’^67^ and related statistics such as the admixture fraction f (also known as the ‘f_4_-ratio) using Dsuite^67, 68^. We focus on the Modern European Newts, which comprise *Triturus*, *Lissotriton*, the NIO clade (*Neurergus*, *Ichthyosaura* & *Ommatotriton*) and *Calotriton*^33^. Calculations are based on the total dataset and on chromosome-level datasets (Table S3).

We conduct the Dsuite analyses for the three different topologies discovered with the phylogenetic analyses earlier: the general species tree based on linkage groups 2-12, with *Lissotriton* being the closest relative to *Triturus*, followed by the NIO clade, then followed by *Calotriton* – named ‘TOP1’), the alternate species tree as inferred with targets from chromosome 1 (with the NIO clade being closer related to *Triturus* than sister lineage *Lissotriton –* named ‘TOP2’), and the deviant species tree as inferred with targets from only 1B (with both *Calotriton* and the NIO clade being more closely related to *Triturus* than sister lineage *Lissotriton –* named ‘TOP3’). The topologies are provided to Dsuite in Newick format. We use *Euproctus* as the outgroup. The Dtrios function of Dsuite is used to calculate the D- and f_4_ admixture ratio statistics for all potential trio combinations. We apply Dsuite’s Fbranch function on trios with significantly elevated D-values to calculate f-branch statistics, which are related to the f4-ratio. Finally, we construct heatmaps using Dsuite as well to visualize the results (Fig. 3, Table S3 and Figs. S36 – S86).

### Determining the number of gene flow events and their direction

The software Treemix allows us to infer maximum likelihood trees and details on admixture events from our target capture data^69, 70^. As input we use the cleaned-up msVCF files that include the Modern European Newt samples with high quality SNPs. We perform more filtering to ensure files have no missing data using VCFtools v.0.1.16^65^ and we remove any sites that are in linkage disequilibrium using PLINK v.1.9^71^. We then run the linkage disequilibrium-pruned data through Treemix, using *Calotriton* as an outgroup with 500 bootstraps, for one through five possible migration edges (m), and for an iteration of ten analyses. Then, we use the R package “OptM”^72^ to determine the optimal value of m using the Treemix output of all iterations. Given that the variation in the standard deviations of the input data of our subsetted datasets appears too low, OptM is not able to retrieve an optimal value for m for smaller subsets of the data (i.e., when including only markers that belong to a certain linkage group), so we decide to only include the overall target capture dataset. We visualize results using (adjusted versions of) the plotting_funcs.R script of Treemix (Fig. 4) and present the Evanno and Linear method outcomes by using the plot_optM function of OptM (Figs. S87 and S88).

### Permit section

*Triturus* embryos were obtained by M.F. from his personal breeding colony. Housing and breeding salamanders as a private individual are not considered to require a license (BGBl. I S. 3125, 3126, 3750). Housing and breeding protocols comply to EU directive standards (EU directive annex III, section B, Table 9.1) and were reviewed by the Animal Welfare Body Leiden. As per EU legislation regarding the protection of animals used for scientific purposes (EU directive no. 2010/63/EU), sacrificing embryos that are not feeding independently does not qualify as an animal experiment. Sacrificing a *Triturus* individual for the Oxford Nanopore sequencing was approved by the Centrale Commissie Dierproeven (CCD; decision no. AVD1060020198065) after advice of the independent Dierexperimentencommissie Leiden (DEC Leiden).

## Acknowledgements

Funding was provided by the European Research Council (ERC) under the European Union’s Horizon 2020 research and innovation programme (Grant Agreement No. 802759). T.B. was supported by DFG (INST 269/768-1). This work was performed using the compute resources from the Academic Leiden Interdisciplinary Cluster Environment (ALICE) provided by Leiden University. We are grateful for the feedback provided to us by Professor Roger K. Butlin from the University of Sheffield, and we thank Erik-Jan Bosch for creating the scientific illustrations of the salamanders under the open content License of Naturalis Biodiversity Center (© CC BY-NC-ND 4.0).

## Author contributions

M.d.V., J.F. and B.W. conceived and designed the research. M.F. collected newt embryos. M.d.V. and J.F. performed the lab-work and pre-processing of the data. M.d.V. conducted the downstream analyses, with the assistance of O.P., C.v.d.P., T.B., and J.F. M.d.V., J.F. and B.W. wrote the draft version of the paper, and all authors contributed to revising it.

## Data Availability

Illumina sequencing reads are openly accessible via the BioProject under PRJNA1173497, PRJNA1171613, and PRJNA498336. Likewise, Oxford Nanopore Technology sequencing reads are available via PRJNA1216568. Scripts for downstream analyses (Dsuite and Treemix) are provided on GitHub (https://github.com/Wielstra-Lab/Triturus_chr1_bloopergenes). Other Supplementary Materials are accessible via Zenodo (https://zenodo.org/records/13991240).

